# Counterfactual MRI Generation with Denoising Diffusion Models for Interpretable Alzheimer’s Disease Effect Detection

**DOI:** 10.1101/2024.02.05.578983

**Authors:** Nikhil J. Dhinagar, Sophia I. Thomopoulos, Emily Laltoo, Paul M. Thompson

## Abstract

Generative AI models have recently achieved mainstream attention with the advent of powerful approaches such as stable diffusion, DALL-E and MidJourney. The underlying breakthrough generative mechanism of denoising diffusion modeling can generate high quality synthetic images and can learn the underlying distribution of complex, high-dimensional data. Recent research has begun to extend these models to medical and specifically neuroimaging data. Typical neuroimaging tasks such as diagnostic classification and predictive modeling often rely on deep learning approaches based on convolutional neural networks (CNNs) and vision transformers (ViTs), with additional steps to help in interpreting the results. In our paper, we train conditional latent diffusion models (LDM) and denoising diffusion probabilistic models (DDPM) to provide insight into Alzheimer’s disease (AD) effects on the brain’s anatomy at the individual level. We first created diffusion models that could generate synthetic MRIs, by training them on real 3D T1-weighted MRI scans, and conditioning the generative process on the clinical diagnosis as a context variable. We conducted experiments to overcome limitations in training dataset size, compute time and memory resources, testing different model sizes, effects of pretraining, training duration, and latent diffusion models. We tested the sampling quality of the disease-conditioned diffusion using metrics to assess realism and diversity of the generated synthetic MRIs. We also evaluated the ability of diffusion models to conditionally sample MRI brains using a 3D CNN-based disease classifier relative to real MRIs. In our experiments, the diffusion models generated synthetic data that helped to train an AD classifier (using only 500 real training scans) -and boosted its performance by over 3% when tested on real MRI scans. Further, we used implicit classifier-free guidance to alter the conditioning of an encoded individual scan to its counterfactual (representing a healthy subject of the same age and sex) while preserving subject-specific image details. From this counterfactual image (where the same person appears healthy), a personalized disease map was generated to identify possible disease effects on the brain. Our approach efficiently generates realistic and diverse synthetic data, and may create interpretable AI-based maps for neuroscience research and clinical diagnostic applications.

## I. Introduction

Magnetic resonance imaging (MRI) scans are collected worldwide for clinical evaluation of disease and for neuroscience research. Alzheimer’s disease (AD) is clinically diagnosed based on neuropsychological and behavioral testing, but imaging data identifies characteristic disease effects on the brain from an early stage [1]. Deep learning methods have also been used for diagnosis, predictive modeling, and dementia subtyping. Convolutional neural networks (CNNs) and vision transformers (ViTs), for example, can be trained to learn from sets of labeled images, where the ground truth diagnosis or patient outcomes are known. These methods usually require additional steps to interpret the model’s decision.

At the same time, several breakthroughs in generative modeling have created novel applications using natural images and medical images. Generative adversarial networks (GANs), for example, are widely used for medical image synthesis [2], image-to-image translation, and super-resolution. The main trade-offs for generative models include fast sampling, diverse and realistic sample generation. GANs can suffer from mode collapse, leading to limited diversity in samples generated and complex training steps. Overcoming this issue, denoising diffusion models have more recently achieved success in generating diverse high-resolution photo-realistic images in both unconditional and conditional (user-guided) contexts [3]. Pinaya et al. trained latent diffusion models (LDM) to generate realistic brain MRI scans, conditioned on demographic information, including age and sex [4]. Others have used variants of DDPMs to synthesize brain MRIs [5] [6] [7] [8]. Diffusion modeling can also be used for counterfactual image generation [9] [10], where an individual image is adjusted to match a reference distribution from healthy controls, while retaining subject-specific anatomical details.

In this work, we trained diffusion models that were conditioned on diagnostic status (here, *AD* and *healthy controls*). We evaluated techniques to train diffusion models with limited data, compute-time and memory resources, by systematically varying the model size, training time, fine-tuning of pre-trained models, and use of a latent diffusion model. We tested the added value of synthetic MRIs - generated by our conditional DDPM - as pre-training data to enhance CNN-based AD classification performance. We generated counterfactual images with the conditional diffusion models with minimal manipulation to preserve subject-specific features in the MRIs. Heat maps were generated from these counterfactual images to visualize possible disease-related abnormalities, offering new feature maps for clinical diagnosis and research.

## II. Data

We used three widely used publicly available brain MRI datasets in our analyses, the UK Biobank [11], ADNI [12] and OASIS [13] (**Table 1**). As in [14], all 3D T1-w brain MRI scans were pre-processed using standard steps for neuroimaging analyses, including: nonparametric intensity normalization (N4 bias field correction) [15], ‘skull-stripping’, linear registration to a template with 9 degrees of freedom, and isotropic resampling of voxels to 2-mm resolution. The input spatial dimension of the MRIs was 91x109x91. All images were scaled to the range 0-1 using min-max scaling, to stabilize model training. The scans were scaled to 32x40x32 for the downscaled DDPM models and cropped to 84x84x84 for the full resolution DDPM models to accommodate computational/memory constraints. The scans were cropped to 80x96x80 for the full-resolution LDM models.

**TABLE I.** Brain T1-w MRI datasets.

| *Dataset* | *Number of Scans* | *Number of Subjects* | *Age range (years)* |
| --- | --- | --- | --- |
| UKB | 38,703 | 38,703 | 44.6-82.8 |
| ADNI | 4,098 | 1,188 | 55.7-92.8 |
| OASIS | 600 | 600 | 43.5-97.0 |

All diffusion models (DMs) were trained or fine-tuned on 3D T1-weighted (T1-w) brain MRI scans from ADNI. We included control samples from the larger UKB dataset, verified based on ICD-10 codes. Additionally, we utilize OASIS as an independent test set to validate our classifier. Relevant demographic information for these datasets is provided in **Table I**.

## III. Methods

The diffusion models are generative in nature and approximate the data distribution using a learned distribution, via a trained neural network. During training, noise sampled from a Normal distribution is added to an image based on a noise scheduler, such as the linear scheduler [4], where noise is added at discrete steps. The diffusion pixel-space based DDPM and LDM models usually consist of a modified U-Net based architecture with a self-attention mechanism. The discrete time steps are also used to *condition* the model and are added to the ResNet blocks of the architecture. The diffusion model is usually designed to predict the added noise at a given random time step, and used to minimize a regression loss function, usually mean squared error (MSE). The trained model is then used to sample synthetic scans via different schedulers, such as DDPM or the deterministic variant denoising diffusion implicit model [16] (DDIM) with a certain number of sampling steps.

During sampling, the model generates data in an iterative manner from pure noise to an image from the training distribution. The choice of the sampling scheduler influences the quality of the synthetic scans and sampling time for each scan. Diffusion models can be conditional if the sampling can further be influenced by additional “context”. Classifier-free (or implicit) guidance [17] involves training the model on both conditional and unconditional data by randomly dropping the class labels (for example, with 15% probability). The conditional and unconditional predictions of the model are then combined via a guidance scale, which offers trade-offs between realism and diversity in the synthetic data.

The conditional version of the DDPM is modified to include a cross-attention mechanism to allow the use of external conditioning using classifier-free guidance such as the “disease” class, i.e., AD or healthy control. We used two main DDPM variants, with a smaller U-Net (4 million parameters) and a larger U-Net (189 million parameters). The MSE loss was the primary loss function for our DDPM models.

We also trained a latent diffusion model (LDM)[3]. LDMs - in contrast to DDPMs - use autoencoders to compress the images, followed by the standard diffusion modeling using the latent space. This step helps alleviate computational bottlenecks that arise when training large deep learning models on large 3D MRI scans. The autoencoder is trained jointly with a reconstruction loss, perceptual loss, KL-loss, and an adversarial loss [4] [3].

## IV. Experiments and Results

### A. Setup

We performed a random search to select hyperparameter values, including the learning rate {1e-3 to 1e-6}, optimizer {ADAM, ADAM with weight decay}, weight decay between 0.1 and 0.0004, batch size {2, 3, 4, 8, 16}. **Table II** summarizes the training hyperparameters. Additionally, during inference for sampling, we used a guidance scale of 2, with 1,000 sampling steps for DDPM, and 250 sampling steps for DDIM.

**TABLE II.**
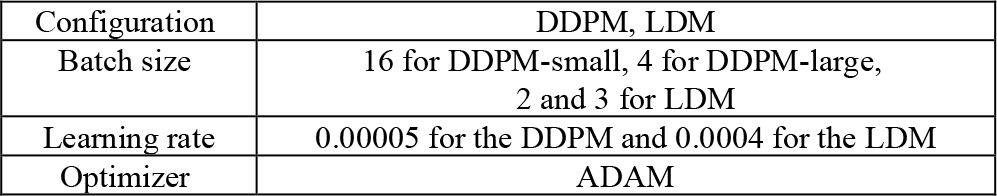
Hyperparameters for model training.

| Configuration | DDPM, LDM |
| --- | --- |
| Batch size | 16 for DDPM-small, 4 for DDPM-large, 2 and 3 for LDM |
| Learning rate | 0.00005 for the DDPM and 0.0004 for the LDM |
| Optimizer | ADAM |

We evaluated our models quantitatively using performance metrics assessing the realism and diversity of the synthetic images. We computed our metrics as average and corresponding standard deviation over 100 evaluation pairs. We used the Maximum Mean Discrepancy (MMD) [18], a kernel based statistical test to measure how similar two distributions are. A smaller MMD value indicates similar distributions. We also used the Multi-scale Structural similarity Metric (MS-SSIM) - another metric to indicate similarity/dissimilarity, where higher values denote greater similarity.

For realism, we evaluated pairs of real images and synthetic images of the same class in each batch of data using MMD and MS-SSIM. Diversity was measured using MS-SSIM comparing two unique synthetic images conditioned on the same class variable, i.e., AD or control. MS-SSIM was also used to measure the similarity in pairs of real images to serve as a reference for diversity in the synthetic data.

### B. Training diffusion models conditioned with disease status

We trained DDPMs and LDMs conditionally on the disease diagnosis (AD, controls) for each subject in ADNI. The LDM training was performed in two stages: the autoencoder was initially trained for 25 epochs, which was followed by training of the diffusion model for 200 epochs. We used two variants for our experiments, a smaller U-Net (4M parameters) and a larger U-Net (189M parameters). We also trained unconditional DDPM and LDM models to serve as a reference for the image quality and diversity of the models.

Further, we evaluated the use of pre-training to overcome limited availability of training data and improve realism, diversity and training efficiency (in terms of compute time and memory). The DDPMs were initially trained to generate T1-w MRI scans conditioned on the “sex” class using the UKBB for pre-training. The pre-training phase was followed by ‘fine-tuning’ involving training on the “disease” class using MRIs from ADNI. The fine-tuning was performed end-to-end. We experimented with training the DDPMs at different image resolutions to test efficiency during training. The data was slightly cropped for training the LDMs, to fit the GPU memory.

## Results

In **Figure 3**. we present synthetic T1-w MRI scans generated using three of the top diffusion-based models. The real MRI scans from ADNI used to train the generative models is shown in **Figure 2** for reference. **Table III** summarizes performance metrics to quantify the synthetic image generation.

**Figure 1.**
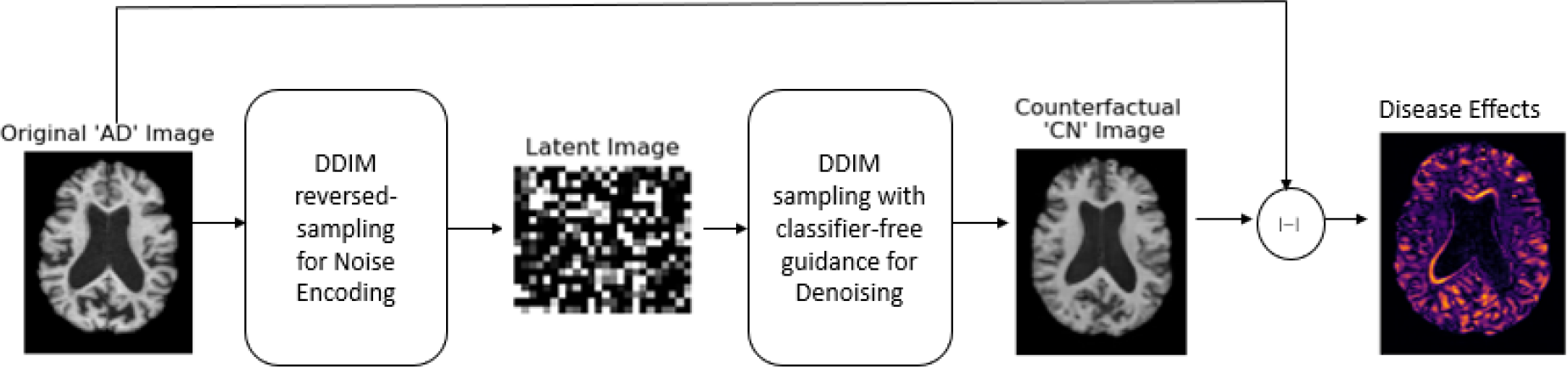
Illustration of the proposed counterfactual image generation framework to identify disease effects in a subject’s T1-w MRI scan.

**Figure 2.**
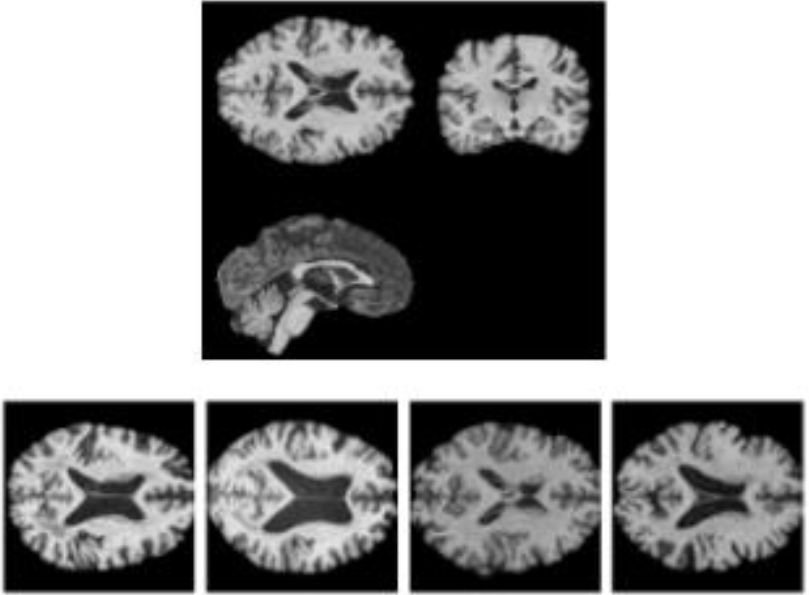
Top - Real T1-w MRI scans from ADNI used for training where each slice is from a different individual. This shows the diversity of the training images, Bottom - Real T1-w MRI scans in 3 planes of section.

**Figure 3.**
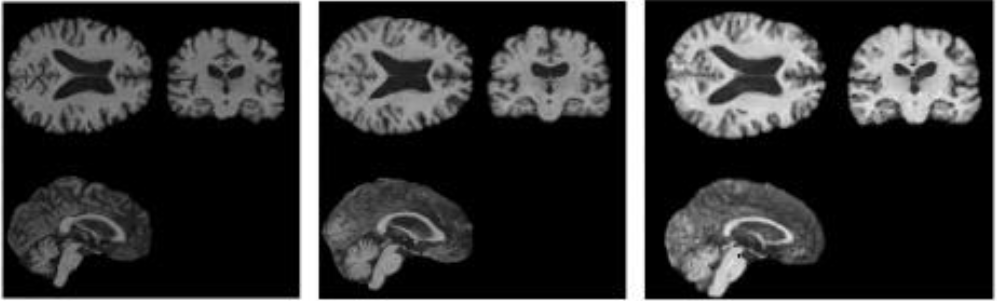
Synthetic T1-w MRIs scans generated with Left - conditional DDPM at full-resolution, Middle - conditional DDPM at full-resolution using pre-training, Right - conditional LDM at full-resolution. The LDM model also uses the additional autoencoder stage during training to compress the images to the latent space, prior to diffusion modeling.

**TABLE III.** Realism and Diversity Test Metrics for the LDMS and pixel-based DDPMS.

| Model (params in millions) | Conditional | Pretrain | Full Resolution | Train epochs | Samp. | Realism: MMD (Real vs Synthetic)↓ | Realism: MS-SSIM (Real vs Synthetic)↑ | Diversity: MS-SSIM (Synthetic vs Synthetic)↓ | Diversity: MS-SSIM (Real vs Real)↓ |
| --- | --- | --- | --- | --- | --- | --- | --- | --- | --- |
| LDM - Small (4M) | - | - | ✓ | 25 (AE) + 200 | DDPM | $0.0182 \pm 0.007$ | $0.847 \pm 0.029$ | $0.869 \pm 0.036$ | $0.841 \pm 0.022$ |
| | | | | | DDIM | $0.019 \pm 0.0086$ | $0.838 \pm 0.035$ | $0.862 \pm 0.045$ | |
| | ✓ | - | ✓ | 25 (AE) + 200 | DDPM | $0.020 \pm 0.010$ | $0.825 \pm 0.048$ | $0.853 \pm 0.063$ | $0.845 \pm 0.025$ |
| | | | | | DDIM | $0.022 \pm 0.012$ | $0.816 \pm 0.054$ | $0.853 \pm 0.068$ | |
| DDPM - Small (4M) | - | - | - | 500 | DDPM | $0.025 \pm 0.019$ | $0.814 \pm 0.107$ | $0.905 \pm 0.049$ | $0.893 \pm 0.028$ |
| | | | | | DDIM | $0.030 \pm 0.019$ | $0.754 \pm 0.109$ | $0.769 \pm 0.106$ | |
| | ✓ | - | - | 500 | DDPM | $0.013 \pm 0.006$ | $0.898 \pm 0.031$ | $0.913 \pm 0.038$ | $0.888 \pm 0.033$ |
| | | | | | DDIM | $0.016 \pm 0.011$ | $0.869 \pm 0.063$ | $0.877 \pm 0.068$ | |
| | ✓ | - | ✓ | 2200 | DDPM | $0.025 \pm 0.013$ | $0.803 \pm 0.057$ | $0.782 \pm 0.129$ | $0.837 \pm 0.029$ |
| | | | | | DDIM | $0.038 \pm 0.021$ | $0.710 \pm 0.127$ | $0.740 \pm 0.212$ | |
| | ✓ | ✓ | ✓ | 500 | DDPM | $0.017 \pm 0.007$ | $0.842 \pm 0.026$ | $0.863 \pm 0.025$ | $0.837 \pm 0.029$ |
| | | | | | DDIM | $0.028 \pm 0.015$ | $0.781 \pm 0.054$ | $0.859 \pm 0.031$ | |
| DDPM - Large (189M) | - | - | - | 500 | DDPM | $0.016 \pm 0.011$ | $0.891 \pm 0.040$ | $0.924 \pm 0.022$ | $0.893 \pm 0.028$ |
| | | | | | DDIM | $0.019 \pm 0.012$ | $0.832 \pm 0.080$ | $0.785 \pm 0.086$ | |
| | ✓ | - | - | 500 | DDPM | $0.013 \pm 0.006$ | $0.893 \pm 0.032$ | $0.908 \pm 0.028$ | $0.888 \pm 0.033$ |
| | | | | | DDIM | $0.019 \pm 0.013$ | $0.834 \pm 0.077$ | $0.832 \pm 0.072$ | |

### C. Evaluation of conditional LDM samples using a 3D CNN for AD Classification

We evaluated the conditional synthetic scans generated by the LDM using a 3D DenseNet121 CNN [19] commonly used as a classifier in neuroimaging research. We trained the CNN on only synthetic data (N = 500 scans), only real data (N = 515 scans), and synthetic data followed by real data (N = 500 + 515 scans). Each of these models was then tested, for AD classification, on an independent test set of 1,219 real MRI T1-w scans from ADNI (not from subjects used in the training set) and 600 real MRI T1-w scans from OASIS for independent validation. The CNN was trained with an initial learning rate of 0.0003 and a finetune learning rate of 0.0001. We used a cosine learning rate scheduler with a warmup of 5 epochs and a binary cross-entropy loss function.

## Results

The evaluation of the synthetic T1-w MRI scans generated by the best performing LDM diffusion model is summarized in **Table IV**. The models tested were either only trained on synthetic data (N = 500 scans), only trained on real data from ADNI (N = 515 scans) or trained initially on synthetic data and then fine-tuned on ADNI (N = 500 + 515 scans).

**TABLE IV.**
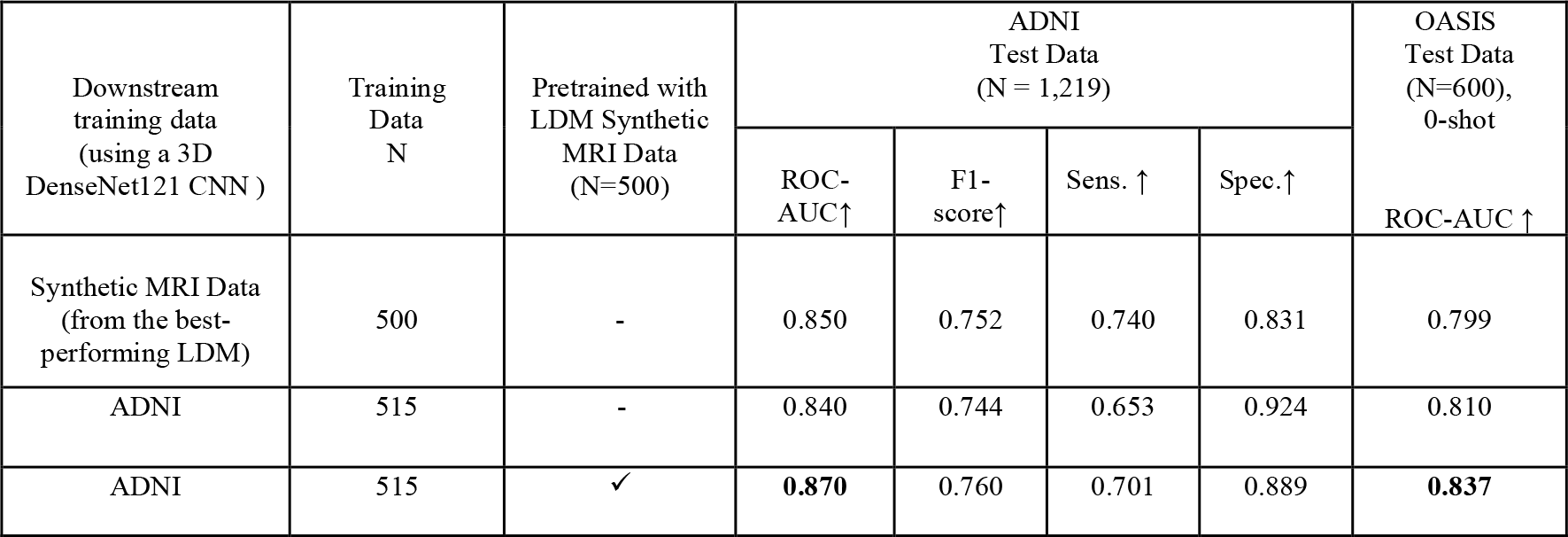
Evaluation of the LDM-generated synthetic T1-w MRI scans via a 3D CNN for AD Classification.

All classification models are tested on real ADNI T1-w MRIs (N = 1,219 scans) and on real OASIS T1-w MRIs (N=600 scans). **Figure 5**. is UMAP of the ADNI test data using the best CNN model’s embedding. Note that in these experiments, the classifier is not used to tell if an image is real or synthetic; the task is to see whether AD classifiers perform better if they are pre-trained on synthetic data.

### D. Counterfactual Image Generation using trained conditional LDMs

We generated counterfactual images by “minimal manipulation” of AD scans to the healthy control distribution during inference [9] [10] using DDIM inversion. DDIM inversion is usually used in image-to-image translation [9] applications as it can maintain aspects of the original image. This mechanism plays an important role in maintaining subject-specific image details during sampling. The sampling process here has two stages, the encoding and decoding stages. The original image is first encoded to the latent space using the trained LDM. The counterfactual is sampled by conditioning the decoding process from the latent. We generated a disease-specific heat map as an absolute difference between the actual AD patient’s scan and the corresponding sampled control counterfactual.

## Results

The control “CN” counterfactual image generated for the T1-w MRI of a subject diagnosed as “AD” in ADNI is shown in **Figure 4**. A counterfactual of “CN” subject from ADNI is also shown as a reference.

**Figure 4.**
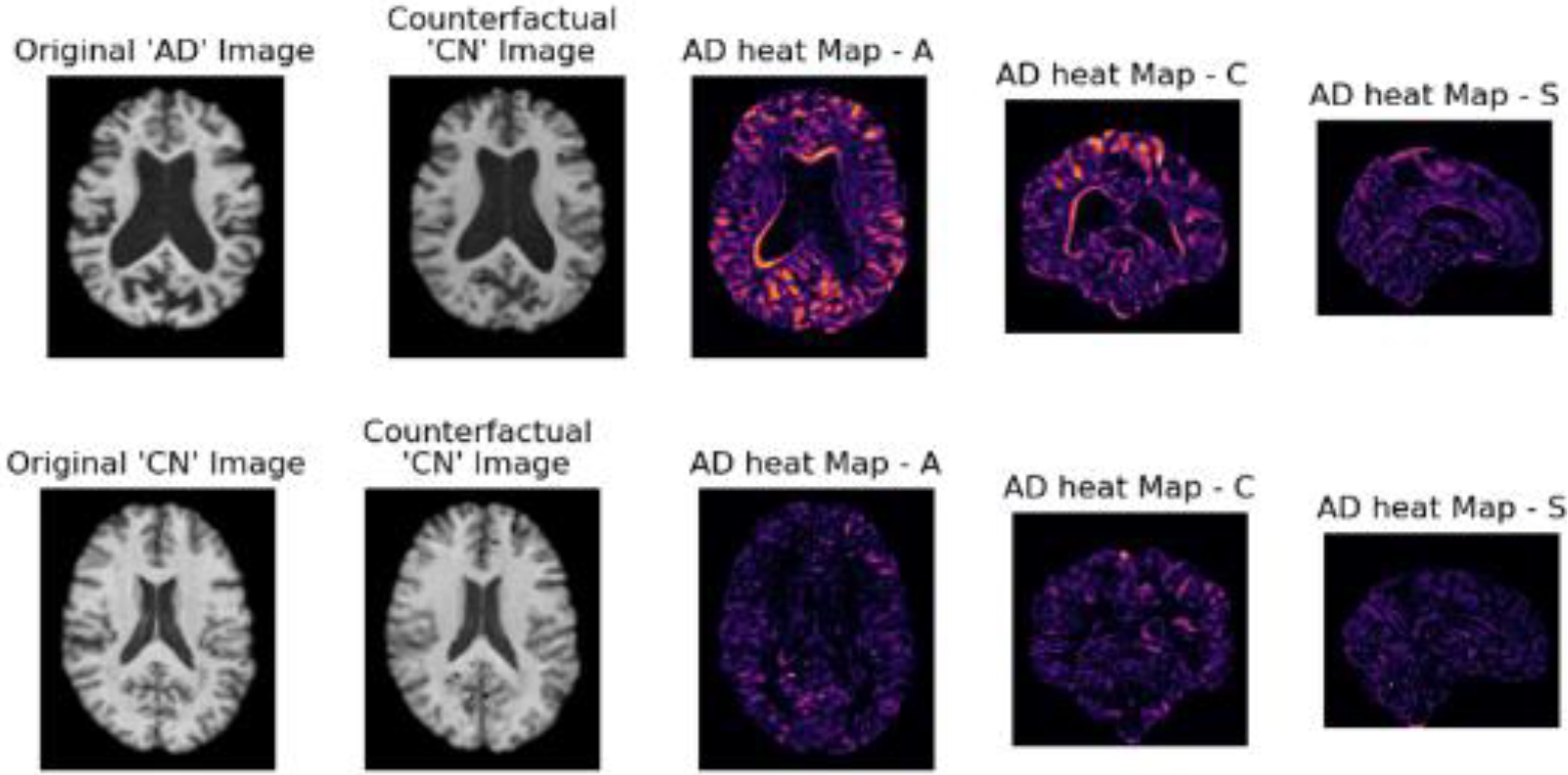
Counterfactual images generated using LDM. Top - T1-w MRI from an actual subject diagnosed as AD from ADNI and the corresponding counterfactual, Bottom - Reference T1-w MRI from a control subject and its reconstruction with minimal changes relative to the AD counterfactual. The disease effects are visualized as a heat map in all three orientations - Axial (A), Coronal (C) and Sagittal (S).

**Figure 5.**
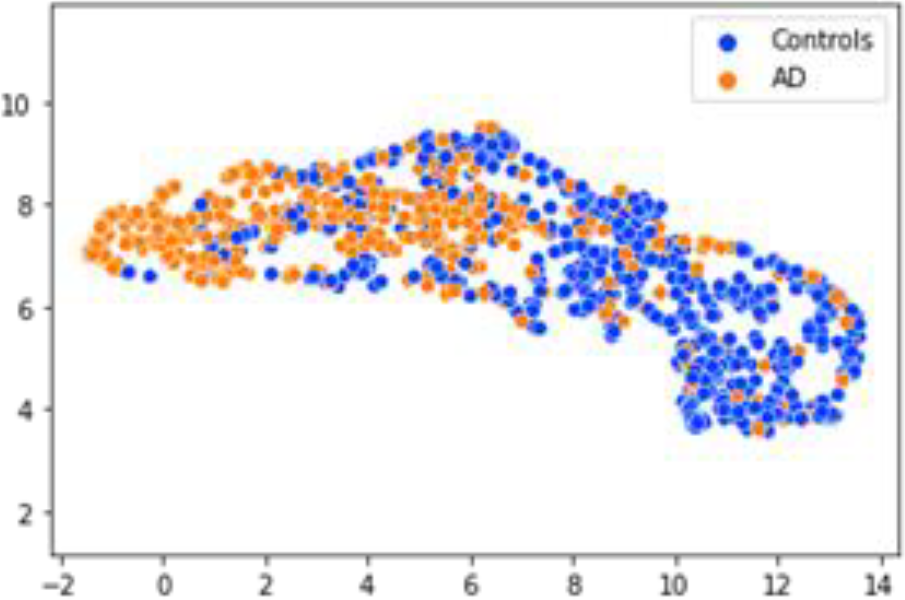
UMAP [20] of class-based clustering with test scans from ADNI when passed through the 3D Densenet121 CNN, first, pre-trained on synthetic LDM MRI data and then fine-tuned.

## V. DISCUSSIONS

### A. Conditional generation in settings with limited training data, compute and memory are benefitted by longer training time, fine-tuning and latent diffusion modelling

In our experiments, LDM-based models provided optimal results when trained from scratch, and were compute-time and memory efficient. We observed that training the DDPM-small model for longer (i.e., with 2,200 epochs), outperformed shorter training runs when using the training data at full resolution. We also identified benefits of fine-tuning pre-trained DDPMs compared to training from scratch. Performance improved with pre-training, yielding comparable results to the LDM. In future work, we will also examine parameter-efficient fine-tuning methods to handle limited downstream training data scenarios.

### B. Synthetic MRIs generated by LDMs can boost AD classification

We evaluated the synthetic MRIs generated by the conditional LDMs, by using them to pre-train a 3D CNN for AD classification. The AD classifier pre-trained on only the LDM-generated MRIs performed similarly, when tested on a held-out dataset of 1,219 real MRIs from ADNI, i.e., with AUC 0.850 vs AUC 0.840. We also found that training with synthetic MRIs and fine-tuning with real MRIs boosted performance by over 3% for AD classification to achieve an 0.870 AUC, which is close to start of the art, using only limited data (as the AD diagnosis itself is not completely reliable, and does not have perfect inter-rater reliability). These experiments were carried out in a limited data setting, using only 515 real and 500 synthetic scans. Additionally using synthetic data to initialize the CNN also boosted performance on 600 real MRI scans from the independent OASIS test dataset when tested in a zero-shot manner i.e., without any additional fine-tuning.

### C. Subject-specific disease maps via counterfactual image generation provides improved interpretability

We generated counterfactual images using DDIM inversion to simultaneously preserve subject-specific image details while converting the image to be close to the healthy control class distribution. Disease heat maps were created as a difference between the real and generated counterfactual images. Such counterfactual images may help provide a basis for a model’s predictions. These heat maps provide additional interpretability and help identify anatomical regions with possible disease effects. Future work will include additional evaluation of these counterfactual images (such as assessing their group statistics) for research and clinical applications.

## VI. Conclusion

In this paper, we tested a counterfactual image generation approach using conditional diffusion models to provide subject-specific interpretation of Alzheimer’s disease in the brain. We trained conditional LDM- and DDPM-based models to generate synthetic 3D T1-weighted MRI scans with the Alzheimer’s disease diagnostic status as the context (conditioning variable). We experimented with variants of the diffusion models in terms of model size, training time and pretraining, to alleviate limitations in the availability of labeled training data.

## References

[1] P. M. Thompson et al., “Tracking Alzheimer’s disease,” Ann. N. Y. Acad. Sci., vol. 1097, pp. 183–214, 2007, doi: 10.1196/annals.1379.017.

[2] G. Kwon, C. Han, and D. shik Kim, “Generation of 3D Brain MRI Using Auto-Encoding Generative Adversarial Networks,” Lect. Notes Comput. Sci. (including Subser. Lect. Notes Artif. Intell. Lect. Notes Bioinformatics), vol. 11766 LNCS, pp. 118–126, 2019, doi: 10.1007/978-3-030-32248-9_14.

[3] R. Rombach, A. Blattmann, D. Lorenz, P. Esser, and B. Ommer, “High-Resolution Image Synthesis with Latent Diffusion Models,” in CVPR, 2022, pp. 10674–10685, doi: 10.1109/cvpr52688.2022.01042.

[4] W. H. L. Pinaya et al., “Brain Imaging Generation with Latent Diffusion Models,” in MICCAI workshop on Deep Generative Models (DGM4MICCAI), 2022, p. pp 117-126, [Online]. Available: http://arxiv.org/abs/2209.07162.

[5] A. Ijishakin, A. Abdulaal, A. Hadjivasiliou, S. Martin, and J. Cole, “Interpretable Alzheimer’s Disease Classification Via a Contrastive Diffusion Autoencoder,” 2023, [Online]. Available: http://arxiv.org/abs/2306.03022.

[6] W. Peng et al., “Metadata-Conditioned Generative Models to Synthesize Anatomically-Plausible 3D Brain MRIs,” pp. 1–26, 2023, [Online]. Available: http://arxiv.org/abs/2310.04630.

[7] Z. Dorjsembe, S. Odonchimed, and F. Xiao, “Three-Dimensional Medical Image Synthesis with Denoising Diffusion Probabilistic Models,” in MIDL, 2022, pp. 2–4, [Online]. Available: https://arxiv.org/abs/2102.09672.

[8] W. Peng, E. Adeli, Q. Zhao, and K. M. Pohl, “Generating Realistic 3D Brain MRIs Using a Conditional Diffusion Probabilistic Model,” 2022.

[9] P. Sanchez, A. Kascenas, X. Liu, A. Q. O’Neil, and S. A. Tsaftaris, “What is Healthy? Generative Counterfactual Diffusion for Lesion Localization,” Lect. Notes Comput. Sci. (including Subser. Lect. Notes Artif. Intell. Lect. Notes Bioinformatics), vol. 13609 LNCS, pp. 34–44, 2022, doi: 10.1007/978-3-031-18576-2_4.

[10] J. Wolleb, F. Bieder, R. Sandkühler, and P. C. Cattin, “Diffusion Models for Medical Anomaly Detection,” Lect. Notes Comput. Sci. (including Subser. Lect. Notes Artif. Intell. Lect. Notes Bioinformatics), vol. 13438 LNCS, pp. 35–45, 2022, doi: 10.1007/978-3-031-16452-1_4.

[11] K. L. Miller et al., “Multimodal population brain imaging in the UK Biobank prospective epidemiological study,” Nat. Neurosci., vol. 19, no. 11, pp. 1523–1536, 2016, doi: 10.1038/nn.4393.

[12] D. P. Veitch et al., “The Alzheimer’s Disease Neuroimaging Initiative in the era of Alzheimer’s disease treatment: A review of ADNI studies from 2021 to 2022,” Alzheimer’s Dement., no. April 2023, pp. 652–694, 2023, doi: 10.1002/alz.13449.

[13] P. J. LaMontagne et al., “OASIS-3: Longitudinal Neuroimaging, Clinical, and Cognitive Dataset for Normal Aging and Alzheimer Disease,” 2019. [Online]. Available: https://www.bertelsmann-stiftung.de/fileadmin/files/BSt/Publikationen/GrauePublikationen/MT_Globalization_Report_2018.pdf http://eprints.lse.ac.uk/43447/1/India_globalisation%2Csocietyandinequalities%28lsero%29.pdf https://www.quora.com/What-is-the.

[14] N. J. Dhinagar et al., “Evaluation of Transfer Learning Methods for Detecting Alzheimer’s Disease with Brain MRI,” 2022.

[15] N. J. Tustison, P. A. Cook, and J. C. Gee, “N4ITK: Improved N3 Bias Correction,” IEEE Trans Med Imaging., vol. 29, no. 6, pp. 1310–1320, 2010, doi: 10.1109/TMI.2010.2046908.N4ITK.

[16] J. Song, C. Meng, and S. Ermon, “Denoising Diffusion Implicit Models,” ICLR 2021 - 9th Int. Conf. Learn. Represent., pp. 1–22, 2021.

[17] J. Ho and T. Salimans, “Classifier-Free Diffusion Guidance,” 2022, [Online]. Available: http://arxiv.org/abs/2207.12598.

[18] A. Gretton, K. M. Borgwardt, M. J. Rasch, B. Schölkopf, and A. Smola, “A kernel two-sample test,” J. Mach. Learn. Res., vol. 13, pp. 723–773, 2012.

[19] G. Huang, Z. Liu, L. van der Maaten, and K. Q. Weinberger, “Densely Connected Convolutional Networks,” in CVPR, 2017, pp. 4700–4708.

[20] L. McInnes, J. Healy, and J. Melville, “UMAP: Uniform Manifold Approximation and Projection for Dimension Reduction,” 2018. [Online]. Available: http://arxiv.org/abs/1802.03426.

